# Distinct lysosomal dysfunction patterns of GRN deficiency in the CNS implicate progranulin in cell type–specific protein sorting

**DOI:** 10.64898/2025.12.29.696915

**Authors:** Gordon C Werthmann, Joachim Herz

## Abstract

Loss-of-function mutations in Progranulin (*GRN*) cause neuronal ceroid lipofuscinosis (NCL) and hereditary frontotemporal dementia, presumably through lysosomal dysfunction. Lysosomes are key metabolic organelles whose functions vary widely depending on their cell type of origin. These functional variations are driven by the lysosomal proteome, yet whether progranulin deficiency alters the lysosomal composition of the mammalian brain in a cell type-specific manner has not been tested. To answer this unknown, we used cell type-specific LysoIP to perform tandem-mass-tag mass-spectrometry and detected distinct aberrant proteomic signatures in progranulin-deficient astrocytes, neurons, and microglia, indicating cell type-specific dysregulation of key lysosomal proteins with crucial functions in sphingolipid metabolism and lysosome organization. These proteins markedly differed from progranulin-deficient RNAseq data sets, suggesting progranulin regulates lysosomal composition through post-translational mechanisms including the sorting of nascent proteins to the lysosome. Validation experiments confirmed that Mfsd8 and Ppt1, proteins whose mutations on their own cause NCL, were essentially absent from progranulin-deficient neuronal and microglial lysosomes, respectively. Our findings demonstrate the protein composition of lysosomes are uniquely sensitive to progranulin deficiency in a cell type-specific manner and that progranulin may function as an essential hub for endolysosomal homeostasis.

## Introduction

Frontotemporal dementia (FTD) is the second most common form of pre-senile dementia and presents with a myriad of symptoms including behavioral abnormalities, impaired executive function, and aphasia (1, 2). Up to 40% of FTD cases are thought to be inherited, and haploinsufficiency of the *GRN* gene on chromosome 17 is one of the most common causes of familial FTD, termed GRN-FTD (3, 4). Patients with GRN-FTD primarily develop the behavioral variant of FTD, and on autopsy, the brains of these patients commonly contain TDP-43 aggregates. *GRN* codes for the protein progranulin, a lysosome resident protein comprised of 7 disulfide bond-rich, full length granulin domains and an N-terminal half-granulin domain. While targeted to the lysosome through various membrane protein interactions (5, 6), progranulin is also known to be secreted into the extracellular space where it is either cleaved into individual granulins or endocytosed and delivered to the lysosome through the endocytic pathway (7, 8).

While haploinsufficiency of *GRN* leads to GRN-FTD, homozygous loss of progranulin leads to a classic lysosomal storage disorder termed CLN11 (neuronal ceroid lipofuscinosis 11) (9). Unlike most proteins associated with lysosomal storage disorders, progranulin or granulins do not appear to have any catalytic function and thus likely function as regulatory proteins within the lysosome. The exact nature of how progranulin regulates the lysosomes, and why loss of progranulin leads to a lysosomal storage disorder are currently unknown, but the links between progranulin and lysosomal function are evident. Loss of progranulin leads to impaired cathepsin D activity (10, 11), aberrant lysosomal lipid composition including decreased levels of the lysosome-specific lipid BMP (bis(monoacylglycerol)phosphate), increased levels of tri-acyl glycerol species (12-14), lysosomal alkalinization (15), and impaired lysosomal protein content *in vitro* (16) and *in vivo (17)*. Progranulin deficiency also activates the CLEAR (coordinated lysosomal expression and regulation) gene network by increasing the nuclear localization of the lysosomal gene master regulator TFEB (transcription factor EB) (18, 19).

Progranulin has further been shown to regulate various cellular functions including neurite outgrowth, immune regulation, and cell migration (20-24). As both a resident lysosomal protein as well as a secreted protein, the mechanism by which progranulin effects such a large range of cellular processes has yet to be fully determined. However, previous work has demonstrated non-secreted, lysosome-targeted progranulin recapitulates the excitotoxic protective effects of wild type progranulin on cultured neurons (25). This suggests the lysosomal function of progranulin is – at least in part – responsible for the non-lysosomal effects of progranulin. Loss of progranulin also leads to a proinflammatory phenotype, and microglia-selective loss of progranulin can recapitulate several phenotypes seen in whole brain progranulin deficiency (26). Progranulin deficient mice demonstrate microgliosis and increased secretion of inflammatory cytokines such as C1qa and C3 (22). *In vitro*, these microglia are sufficient to induce aggregation of TDP-43 in co-cultured neurons (27).

How progranulin regulates lysosomal function remains to be understood; however, previous work has demonstrated loss of progranulin leads to aberrations of the lysosomal proteome, dysregulating key lysosomal proteins that are also implicated in lysosomal storage disorders (16, 17). Lysosomes take on unique functions depending on their cell type of origin (28, 29), and previous work has demonstrated the cell types of the mammalian brain also have unique lysosomal proteomes (30). We hypothesized that lysosomal proteomes are uniquely sensitive to progranulin in a cell type-dependent manner. In order to test this, we utilized a technology termed LysoTag which allows for the isolation of lysosomes *in vivo* in a cell type-specific manner in order to investigate the effects of progranulin deficiency on the lysosomal proteome (31). The specificity of this technique allowed for precise isolation and investigation of the lysosomal proteome in progranulin-deficient neurons, microglia, and astrocytes, revealing cell-type dependent protein aberrations not evident upon whole brain analysis nor at the transcript level, and demonstrating the ability of progranulin to regulate cell-specific lysosomal protein homeostasis in the mammalian brain.

## Results

### LysoIP isolates lysosomes from specific cell types in *Grn* WT, Het, and KO mice

In order to investigate the effects of progranulin deficiency on the lysosomal proteome *in vivo*, we bred four specific Cre recombinase lines with previously defined TMEM192-3xHA fl/fl (30, 31) and *Grn* KO mouse lines (13). These included *Ubc-*, *Syn-*, *Cx3cr1-*, and *Aldh1l1-Cre* lines for pancellular, neuron-, microglia-, and astrocyte-specific LysoIP, respectively (Figure 1a). LysoIP was performed on the brains of 2 to 3-month-old *Ubc-*, *Syn-*, and *Cx3cr1-Cre* mice or 6-month-old *Aldh1l1-Cre* mice. 6-month-old *Aldh1l1-Cre* mice were used, because progranulin was undetectable in any 3-month-old *Aldh1l1-Cre* mouse LysoIP sample including those derived from *Grn* WT mice. This is in line with progranulin being primarily expressed in neurons and microglia (32). Protein concentrations were normalized prior to tandem mass tag mass spectrometry (TMT-MS). Splenic macrophage-derived lysosomes were also obtained from *Cx3cr1-Cre* mice to further characterize the effect of progranulin loss on immune cell lysosomes. Western blot verification demonstrated reliable and specific isolation of lysosomes from all four Cre lines (Figure 1b-e). Interestingly, in *Cx3cr1-Cre* LysoIP samples, Lamp1, a marker for lysosomal membranes, demonstrated a molecular weight shift from ∼75 kDa to ∼110 kDa (Figure 1c). This was seen in both whole-cell lysate and LysoIP samples from mouse spleens – an organ primarily comprised of immune cells (Supplemental Figure 1a), suggesting this shift is specific to immune cells and is thus unlikely to be seen in whole-brain lysates. Lamp1 has various glycosylation sites which likely lead to this mass shift (33).

**Figure 1:**
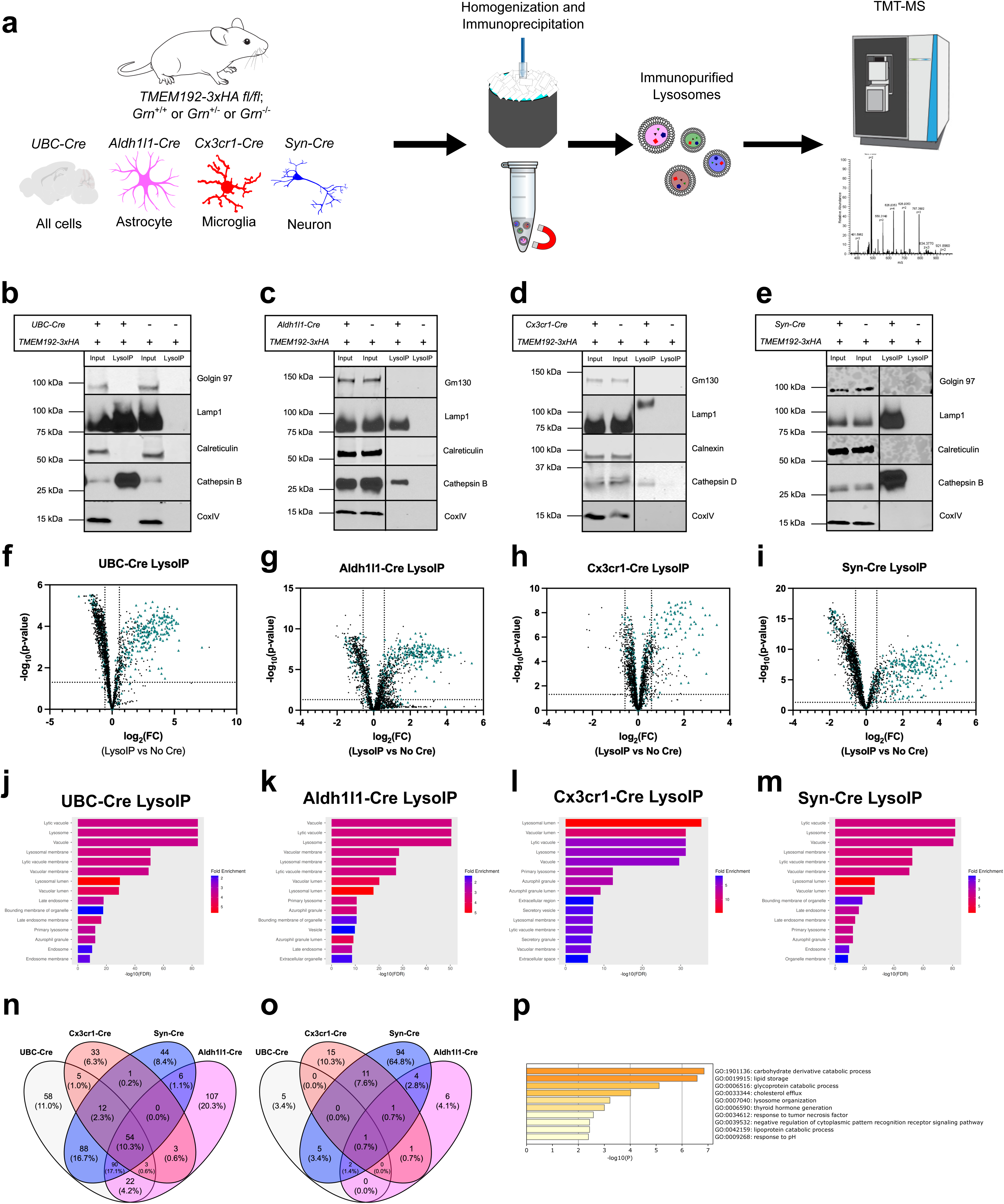
*In vivo* LysoIP isolates cell-type-specific lysosomes from progranulin deficient mice. **a** Experimental design for cell-type-specific lysosomal isolation from progranulin deficient mice followed by TMT-MS. **b-e** Western blot verification of *in vivo* LysoIP from all brain cells (*Ubc-Cre*, **b**), microglia (*Cx3cr1-Cre*, **c**), astrocytes (*Aldh1l1-Cre*, **d**), and neurons (*Syn-Cre*, **e**). **f-i** Volcano plots comparing LysoIP samples to no *Cre* controls (adjusted p-value < 0.05, one-sample t-test followed by Benjamini-Hochberg correction). Known lysosomal proteins are marked as blue triangles. **j-m** Gene ontology analysis of LysoIP samples from four separate *Cre* lines after processing of raw TMT-MS peak intensities. **n** Venn diagram that demonstrates overlap of all brain-derived, lysosomal proteins found in four separate *Cre* lines. **o** Venn diagram that demonstrates overlap of significantly altered brain-derived, lysosomal proteins (*Grn* WT vs *Grn* KO, adjusted p-value < 0.05) in four separate *Cre* lines. **p** Gene ontology analysis of significantly altered, brain-derived lysosomal proteins found in all four *Cre* lines (*Grn* WT vs *Grn* KO, adjusted p-value < 0.05).

After TMT-MS, raw protein intensities were processed and normalized as described in Methods (Supplemental Figure 1b-e). Several thousand proteins were detected in each TMT-MS run (*Ubc-Cre*: 2310 proteins, *Cx3cr1-Cre* (Brain): 2302 proteins, *Cx3cr1-Cre* (Spleen): 1972 proteins, *Aldh1l1-Cre*: 2263 proteins, and *Syn-Cre*: 2219 proteins). To ensure stringent analysis of the lysosomal proteome only, LysoIP samples were compared to no *Cre* control samples added to each TMT-MS run (Figure 1f-i, Supplemental Figure 1f), and proteins were only included for further analysis if they were significantly enriched in LysoIP samples compared to no *Cre* controls and if their normalized abundance was at least 1.5x that of no *Cre* controls. This resulted in hundreds of proteins per *Cre* line (*Ubc-Cre*: 332 proteins, *Cx3cr1-Cre* (Brain): 111 proteins, *Cx3cr1-Cre* (Spleen): 147 proteins, *Aldh1l1-Cre*: 285 proteins, and *Syn-Cre*: 295 proteins). Gene ontology analysis of the resultant proteins demonstrated significant enrichment for lysosomal compartments (Figure 1j-m).

Between all TMT-MS runs, 526 unique lysosomal proteins were detected in both *Grn* WT and *Grn* KO LysoIP samples. Several of these proteins were shared between at least 2 individual Cre lines; however, 45.4% of all proteins detected were only detected in a single Cre line (Figure 1f). This most likely represents a cell type-specific lysosomal proteomic landscape(34). These proteins were then compared between *Grn* WT, *Grn* heterozygous, and *Grn* KO LysoIP samples of the same Cre using ANOVA and multiple comparison correction as outlined in Methods. Any protein with an adjusted p-value < 0.05 was considered significant. Comparing significantly altered proteins between Cre lines, most significantly altered proteins were detected in neuronal lysosomes (*Syn-Cre*) (Figure 1o, j). When comparing the number of significantly altered proteins in individual Cre lines to the total number of proteins detected, microglial and neuronal lysosomes demonstrated the highest percentage of significantly altered proteins (26.1% and 40.0%, respectively) (Supplemental Figure 1h-k). This is consistent with progranulin primarily being expressed in microglia and neurons. Gene ontology analysis of all significantly altered lysosomal proteins across Cre lines – using all 526 lysosomal proteins as background – highlights carbohydrate derivative catabolism, lipid storage, lysosomal organization, glycoprotein catabolism, and cholesterol efflux as some of the most highly enriched cellular processes affected by progranulin deficiency (Figure 1p; Supplemental Table 1).

### Changes to progranulin-deficient lysosomes are not reflected in mRNA or whole-cell lysate changes

Comparing *Ubc-Cre*+ *Grn* WT and *Grn* KO samples, progranulin deficiency significantly altered 13/332 proteins with 8 of these proteins being upregulated and 5 downregulated (Figure 2a-c; Supplemental Table 2) while lysosomes from *Grn* heterozygous mouse brains did not demonstrate any significant changes (Figure 2b). Proteins associated with endo-lysosomal trafficking such as Tspan7 were significantly upregulated in *Grn* KO *Ubc-Cre* LysoIP samples (35, 36). Tspan 7 has also been associated with synaptic transmission(37, 38). Interestingly, Tthy2 and Ttyh3 are also significantly upregulated in *Grn* KO *Ubc-Cre* LysoIP samples. Tthy2 was recently shown to interact with ApoE and facilitate endosomal lipid transfer (39). The APOE4 allele is the strongest genetic predictor for Alzheimer’s disease (AD), and patients with GRN mutations are also genetically predisposed to AD, suggesting the effect of progranulin deficiency on lysosomal Tthy2 may contribute to this increased risk of developing AD (40, 41).

**Figure 2.**
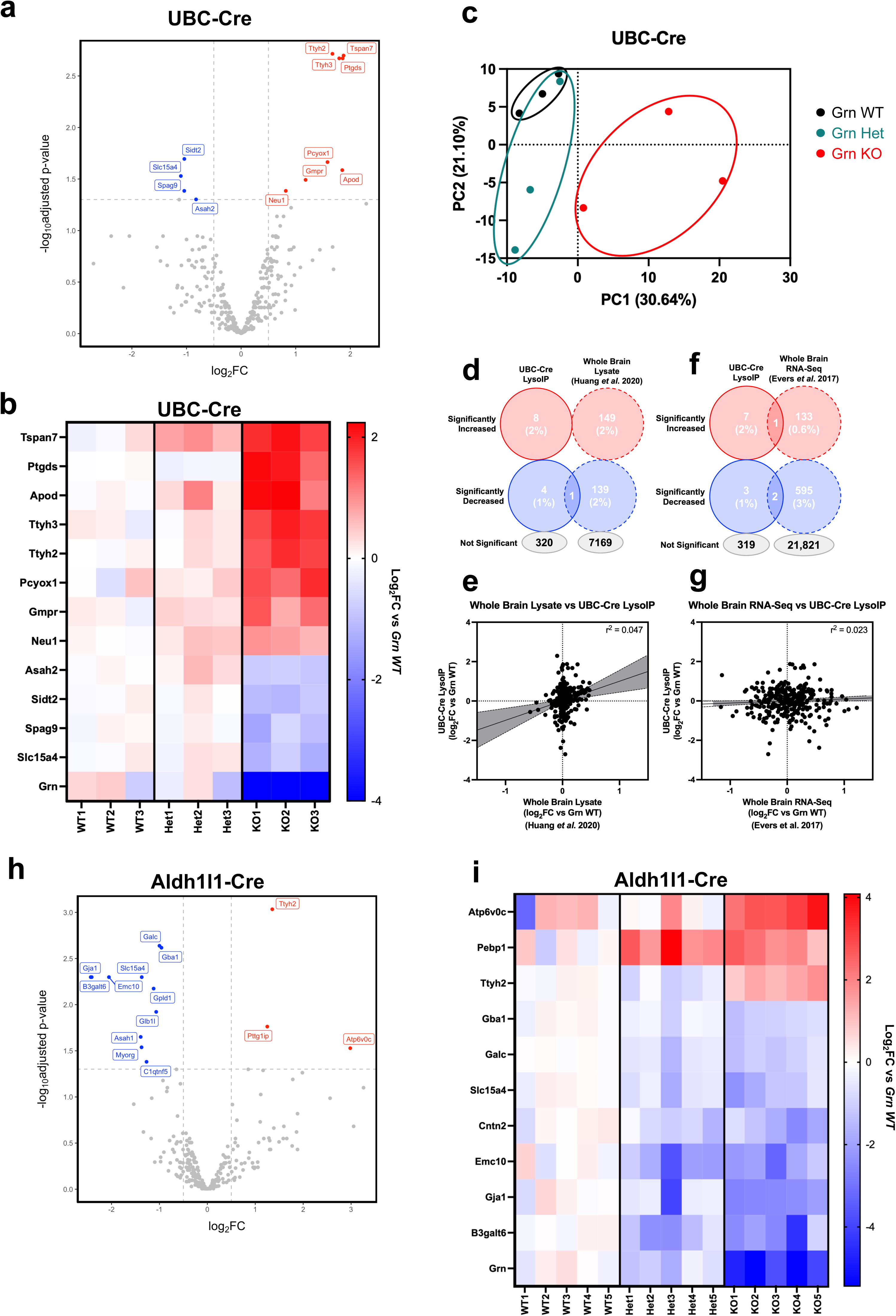
*In vivo* LysoIP analysis of 3-month-old *Ubc-Cre* mouse brains demonstrates post-translational alterations in key lysosomal proteins. **a** Volcano plot demonstrating significantly altered proteins (*Grn* WT vs *Grn* KO adjusted p-value < 0.05) from whole-brain LysoIP TMT-MS samples (n = 3 *Grn* WT, n = 3 *Grn* Het, n = 3 *Grn* KO). **b** Principal component analysis comparing log_2_FC values between *Grn* WT, *Grn* Het, and *Grn* KO brain-derived, lysosomal proteins in *Ubc-Cre* mice. **c** Heatmap demonstrating log_2_FC values for significantly altered, brain-derived lysosomal proteins found in brain-derived *Ubc-Cre* LysoIP TMT-MS samples (*Grn* WT vs *Grn* KO, adjusted p-value < 0.05). **d** Venn diagram of the overlap of significantly altered proteins (*Grn* WT vs *Grn* KO, adjusted p-value < 0.05) between brain-derived *Ubc-Cre* LysoIP and previously published whole brain, whole cell lysate TMT-MS experiments. **e** Venn diagram of the overlap of significantly altered proteins/transcripts (*Grn* WT vs *Grn* KO, adjusted p-value/q-value < 0.05) between brain-derived *Ubc-Cre* LysoIP TMT-MS samples and previously published whole brain RNA-Seq data. **f** Comparison of all log_2_FC values (*Grn* WT vs *Grn* KO) for proteins found in both brain-derived *Ubc*-*Cre* LysoIP TMT-MS samples and previously published whole brain TMT-MS datasets **g** Comparison of all log_2_FC values (*Grn* WT vs *Grn* KO) for proteins/transcripts found in both brain-derived *Ubc*-*Cre* LysoIP TMT-MS samples and previously published whole brain RNA-Seq datasets. **h** Volcano plot demonstrating significantly altered proteins (*Grn* WT vs *Grn* KO adjusted p-value < 0.05) from whole-brain *Aldh1l1-Cre* LysoIP TMT-MS samples (n = 5 *Grn* WT, n = 5 *Grn* Het, n = 5 *Grn* KO). **i** Heatmap demonstrating log_2_FC values for significantly altered, brain-derived lysosomal proteins found in brain-derived *Aldh1l1-Cre* LysoIP TMT-MS samples (*Grn* WT vs *Grn* KO, adjusted p-value < 0.05).

We next compared progranulin-deficient LysoIP samples against TMT-MS results of progranulin-deficient whole brain lysates (Figure 2d-e) (42). 237 proteins were shared between these datasets; however, only progranulin was found to be significantly altered in both LysoIP and whole brain samples. There was overall poor correlation between progranulin-dependent changes from these datasets, suggesting the effects of progranulin deficiency on the lysosomal proteome can primarily only be detected by directly measuring lysosomal proteins as previously reported (43). Further, comparisons between log_2_FC’s of the 301 proteins detected in both our *Grn* KO LysoIP samples and *Grn* KO RNA-Seq datasets from Evers *et al.* 2017 similarly demonstrate lack of overlap in significantly altered proteins (Figure 2f) (13), and when comparing all proteins detected in *Ubc-Cre* LysoIP samples to their respective mRNA transcripts from whole-brain RNA-Seq data, no significant correlation can is detected (Figure 2g). This implies progranulin deficiency alters the lysosomal proteome at the post-transcriptional level.

### Progranulin deficiency causes minor changes in astrocyte lysosomes

Progranulin expression in astrocytes is relatively low compared to other cell types in the brain (32). In fact, TMT-MS analysis of 3-month-old *Aldh1l1-Cre* LysoIP samples did not detect progranulin. However, progranulin was detected in 6-month-old *Aldh1l1-Cre* LysoIP samples, and comparisons of these samples demonstrated 15/285 significantly altered lysosomal proteins between *Grn* WT vs *Grn* KO mouse brains, with 3 being significantly upregulated and 12 being significantly downregulated (Figure 2h-i, Supplemental Table 3). Similar to *Ubc-Cre* experiments, the membrane protein Ttyh2 is upregulated in *Grn* KO astrocyte lysosomes, and several proteins involved in sphingolipid metabolism such as Galc and Gba1 are downregulated (44, 45). Galc and Gba1 mutations have also both been implicated in Parkinsons disease (46, 47).

### Progranulin deficiency dysregulates various disease-related proteins in the neuronal lysosome

In contrast to the minimal effect of progranulin loss on *Ubc-Cre+* and *Aldh1l1-Cre*+ LysoIP samples, progranulin deficiency heavily influences the neuronal lysosomal proteome, leading to 118/295 proteins being significantly altered with 59 proteins being upregulated and 59 being downregulated (Figure 3a-b; Supplemental Table 4). PCA demonstrates clear delineation between *Grn* WT and *Grn* KO neuronal lysosomes while *Grn* heterozygous lysosomes are more heterogeneous between samples (Figure 3c). When investigating the cellular functions related to significantly altered proteins in *Grn* KO neuronal lysosomes, gene ontology analysis demonstrates sphingolipid biosynthesis, lipid biosynthesis, and lysosome organization as the most highly enriched terms (Figure 3d; Supplemental Table 5). Proteins associated with ceramide metabolism such as Neu4, Asah1, Asah2, and Gba1 are all downregulated in *Grn* KO neuronal lysosomes (Figure 3e). Key autophagy proteins that are part of mTORC1 including mTor, Mlst8, and Rptor are all significantly upregulated in *Grn* KO neuronal lysosomes while the adaptor proteins Lamtor2, Lamtor4, and the amino acid transporter Slc38a9 are all significantly downregulated(48). This dysregulation may contribute to the impaired autophagy previously described in models of progranulin deficiency (49, 50).

**Figure 3.**
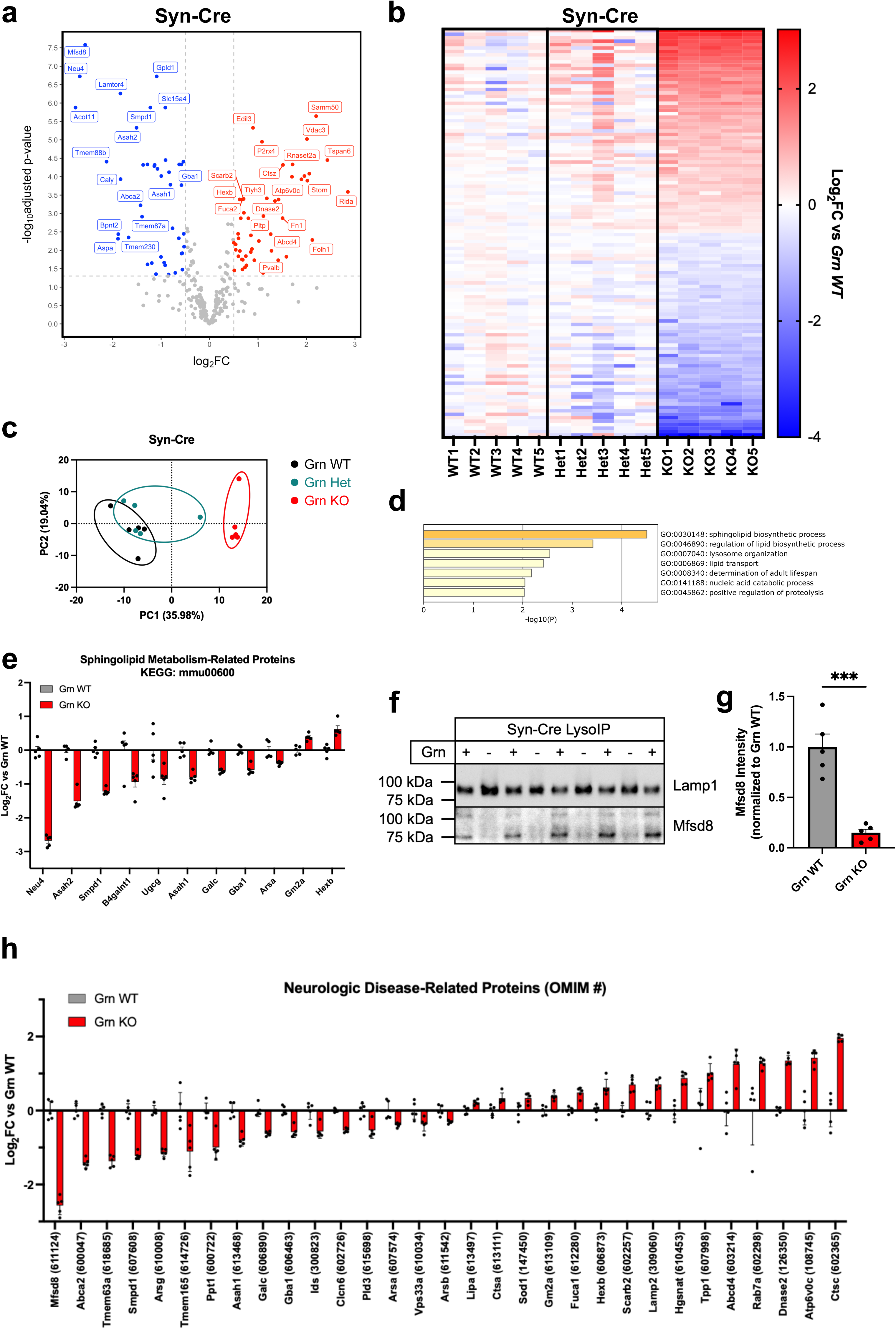
*In vivo* LysoIP analysis of 3-month-old *Syn-Cre* mouse brains demonstrates aberrations in disease-relevant lysosomal proteins in progranulin-deficient neurons. **a** Volcano plot demonstrating significantly altered proteins (*Grn* WT vs *Grn* KO adjusted p-value < 0.05) from neuronal LysoIP TMT-MS samples (n = 5 *Grn* WT, n = 5 *Grn* Het, n = 5 *Grn* KO). **b** Heatmap demonstrating log_2_FC values for significantly altered neuronal LysoIP samples (*Grn* WT vs *Grn* KO, adjusted p-value < 0.05). **c** Principal component analysis comparing log_2_FC values in neuronal lysosomal proteins between *Grn* WT, *Grn* Het, and *Grn* KO mice. **d** Gene ontology analysis of significantly altered neuronal lysosomal proteins (*Grn* WT vs *Grn* KO, adjusted p-value < 0.05). **e** Significantly altered neuronal lysosomal proteins (*Grn* WT vs *Grn* KO, adjusted p-value < 0.05) which are associated with sphingolipid metabolic processes based on gene ontology analysis. **f-g** Western blot (**f**) and densitometry analysis (**g**) of Mfsd8 in neuronal lysosomes (n = 5 *Grn* WT, n = 4 *Grn* KO; ***p < 0.001, Students’ t-test) **h** Significantly altered neuronal lysosomal proteins (*Grn* WT vs *Grn* KO, adjusted p-value < 0.05) whose mutations are associated with neurologic disease.

Several mitochondrial proteins including Vdac1/2/3 and Samm50 are all upregulated in *Grn* KO neuronal lysosomes (Supplemental Figure 2a) (51-54). Progranulin deficiency has been implicated in mitophagy defects(55, 56), so we performed Seahorse analysis to determine if mitochondria in *Grn* KO mouse brains demonstrated decreased function. We did not observe any overt mitochondrial deficits using this method (Supplemental Figure 2b-c). However, we did detect a significantly decreased NAD+/NADH ratio from *Grn* KO mouse brains compared to *Grn* WT (Supplemental Figure 2d-f). This was driven by both an increase in NADH and a decrease in NAD+ in *Grn* KO mouse brains and demonstrates how progranulin deficiency alters brain energy homeostasis.

Mfsd8 was the most significantly downregulated protein between *Grn* WT and *Grn* KO LysoIP samples. Homozygous MFSD8 nonsense mutations lead to a classic lysosomal storage disorder (CLN7), and missense mutations have also been found to be associated with FTD in a small subset of patients (57). Western blot analysis verified Mfsd8 was heavily decreased in *Grn* KO neuronal lysosomes (Figure 3f-g). Mfsd8 was not detected in either astrocyte or microglial LysoIP samples, suggesting it is specific to neuronal lysosomes and thus only regulated by neuronal progranulin.

Of the proteins significantly dysregulated in *Grn* KO neuronal lysosomes, 32 of them are known lysosomal proteins associated with known neurologic disorders based on the OMIM database (Figure 3h, Table 1) (34). This includes proteins associated with lysosomal storage disorders such as Mfsd8, Smpd1, Galc, and Asah1 (44, 58-60).

**Table 1:** List of endo-lysosomal disease related proteins significantly altered in progranulin-deficient neuronal lysosomes.

| Gene Name | OMIM# | Disease | Log<sub>2</sub>FC<br>(*Grn* KO/*Grn* WT) |
| --- | --- | --- | --- |
| Abca2 | 600047 | Intellectual developmental disorder with poor growth and with or without seizures or ataxia | -1.422 |
| Abcd4 | 603214 | Methylmalonic aciduria and homocystinuria, cblJ type | 1.2586 |
| Arsa | 607574 | Metachromatic leukodystrophy | -0.3898 |
| Arsb | 611542 | Mucopolysaccharidosis type VI (Maroteaux-Lamy) | -0.3096 |
| Arsg | 610008 | Usher syndrome, type IV | -1.14 |
| Asah1 | 613468 | Farber lipogranulomatosis | -0.8072 |
| Atp6v0c | 108745 | Epilepsy, early-onset, 3, with or without developmental delay | 1.424 |
| Clcn6 | 602726 | Ceroid lipofuscinosis, neuronal, 15 | -0.5394 |
| Ctsa | 613111 | Galactosialidosis | 0.3282 |
| Ctsc | 602365 | Haim-Munk syndrome | 1.958 |
| Dnase2 | 126350 | Autoinflammatory-pancytopenia syndrome | 1.35 |
| Fuca1 | 612280 | Fucosidosis | 0.4864 |
| Galc | 606890 | Krabbe disease | -0.6264 |
| Gba1 | 606463 | Gaucher Disease | -0.5804 |
| Gm2a | 613109 | GM2-gangliosidosis, AB variant | 0.3804 |
| Grn | 138945 | Ceroid lipofuscinosis, neuronal, 11 | -4.226 |
| Hexb | 606873 | Sandhoff disease, infantile, juvenile, and adult forms | 0.6222 |
| Hgsnat | 610453 | Mucopolysaccharidosis type IIIC (Sanfilippo C) | 0.8754 |
| Ids | 300823 | Mucopolysaccharidosis II | -0.5634 |
| Lamp2 | 309060 | Danon disease | 0.7078 |
| Lipa | 613497 | Cholesteryl ester storage disease | 0.2158 |
| Mfsd8 | 611124 | Ceroid lipofuscinosis, neuronal, 7 | -2.568 |
| Pld3 | 615698 | Spinocerebellar ataxia 46 | -0.5342 |
| Ppt1 | 600722 | Ceroid lipofuscinosis, neuronal, 1 | -0.9972 |
| Rab7a | 602298 | Charcot-Marie-Tooth disease, type 2B | 1.28 |
| Scarb2 | 602257 | Epilepsy, progressive myoclonic 4, with or without renal failure | 0.7028 |
| Smpd1 | 607608 | Niemann-Pick disease, type A/B | -1.222 |
| Sod1 | 147450 | Parkinson's Disease | 0.3346 |
| Tmem165 | 614726 | Congenital disorder of glycosylation, type IIk | -1.1058 |
| Tmem63a | 618685 | Leukodystrophy, hypomyelinating, 19, transient infantile | -1.368 |
| Tpp1 | 607998 | Ceroid lipofuscinosis, neuronal, 2 | 1.0158 |
| Vps33a | 610034 | Mucopolysaccharidosis-plus syndrome | -0.371 |

### Progranulin deficiency dysregulates disease-relevant proteins in the microglial lysosome

While fewer total lysosomal proteins were detected in *Cx3cr1-Cre* LysoIP samples – likely due to less starting material – several key lysosomal proteins were found to be dysregulated secondary to progranulin deficiency with a total of 29/111 proteins being significantly altered with 14 proteins being upregulated and 15 being downregulated (Figure 4a-c; Supplemental Table 6). Based on GO analysis, progranulin deficiency in microglia primarily affects lysosomal proteins associated with lipid catabolism and tissue regeneration, both processes known to be dysregulated in models of progranulin deficiency (13, 61) (Figure 4d; Supplemental Table 7).

**Figure 4.**
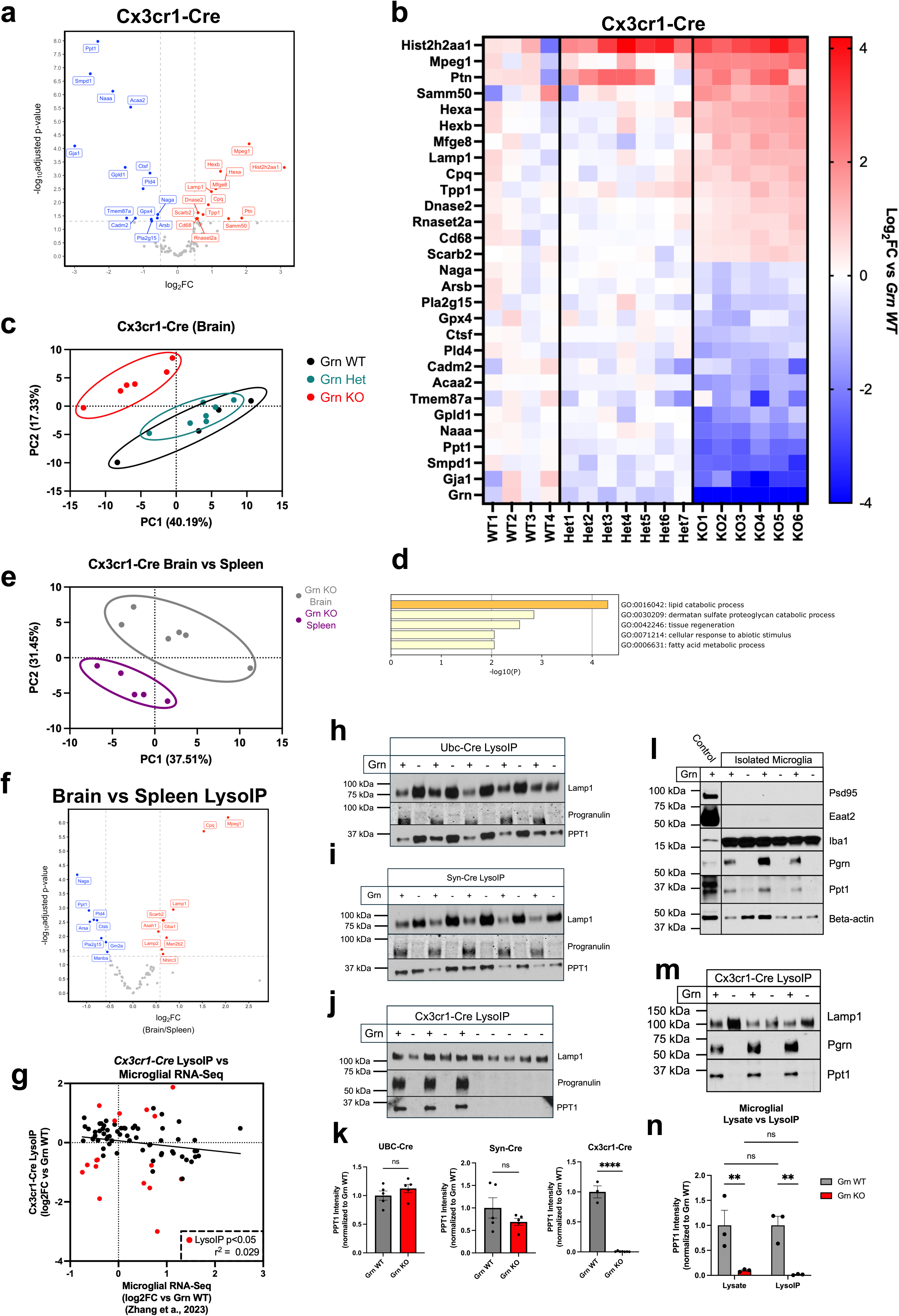
*In vivo* LysoIP analysis of 3-month-old *Cx3cr1-Cre* mouse brains demonstrates cell-specific loss of lysosomal storage disease-related protein in progranulin-deficient microglia. **a** Volcano plot demonstrating significantly altered proteins (*Grn* WT vs *Grn* KO adjusted p-value < 0.05) from microglial LysoIP TMT-MS samples (n = 4 *Grn* WT, n = 7 *Grn* Het, n = 6 *Grn* KO). **b** Gene ontology analysis of significantly altered microglial lysosomal proteins (*Grn* WT vs *Grn* KO, adjusted p-value < 0.05). **c** Principal component analysis comparing log_2_FC values in microglial lysosomal proteins between *Grn* WT, *Grn* Het, and *Grn* KO mice. **d** Heatmap demonstrating log_2_FC values for significantly altered microglial LysoIP samples (*Grn* WT vs *Grn* KO, adjusted p-value < 0.05). **e** Principal component analysis comparing log_2_FC (*Grn* WT vs *Grn* KO) between brain- and spleen-derived *Cx3cr1-Cre* LysoIP samples. **f** Volcano plot demonstrating significantly altered proteins (brain vs spleen adjusted p-value < 0.05) from brain-and spleen-derived *Cx3cr1-Cre* LysoIP TMT-MS samples (n = 6 brain-derived, n = 6 spleen-derived). **g** Comparison of log2FC values (*Grn* WT vs *Grn* KO) between *Cx3cr1-Cre* LysoIP samples and microglial mRNA transcripts from a previously published RNA-Seq dataset. Red circles indicate proteins significantly altered in *Cx3cr1-Cre+*;*Grn* KO LysoIP samples. **h-j** Western blot analysis of Ppt1 in whole-brain (**h**) neuronal (**i**), and microglial (**j**) lysosomes derived from *Grn* WT and KO mouse brains. **k** Densitometry analysis of Ppt1 protein in whole-brain, neuronal, and microglial lysosomes derived from *Grn* WT and KO mouse brains. Values were standardized to Lamp1 and normalized to *Grn* WT (whole-brain: n = 5 3-month-old *Grn* WT and n = 5 3-month-old *Grn* KO; neuronal: n = 5 3-month-old *Grn* WT and n = 5 3-month-old *Grn* KO; microglia: n = 3 6-month-old *Grn* WT and n = 7 6-month-old *Grn* KO; ****p < 0.0001, Students’ t-test). **l-m** Western blot analysis of Ppt1 in microglial lysosomes (**l**) and whole microglia (**m**). Psd95, Eeat2, and Iba1 act as neuronal, astrocyte, and microglial markers, respectively. **n** Densitometry analysis of Ppt1 blots. Values were standardized to Lamp1 (**l**) or beta-actin (**m**) and normalized to *Grn* WT (n = 3 9-12-month-old *Grn* WT and n = 3 9-12-month-old *Grn* KO; **p < 0.01, two-way ANOVA with Fisher’s LSD test).

To test whether the effects of progranulin deficiency on lysosomal protein composition were specific to microglia or were universal to all immune cells, we performed TMT-MS analysis on LysoIP samples derived from spleens of *Cx3cr1-Cre*+; *Grn* WT and KO mice. This analysis revealed several proteins significantly altered in *Grn* KO splenic macrophage lysosomes compared to *Grn* WT (Supplemental Figure 3a-c; Supplemental Table 8). However, when comparing proteins between *Grn* KO microglial and splenic macrophages by principal component analysis, the two tissue types clustered separately (Figure 4e) and several proteins were found to be significantly different (Figure 4f). This suggests the effects of progranulin deficiency on immune cells are in part dependent on the tissue of origin. Of note, Pld4 is significantly downregulated in progranulin deficient microglia (log_2_FC = -1.00, adjusted p-value = 0.003). Pld4 was recently discovered to be a synthetic protein for BMPs within the lysosomes of immune cells (62). BMPs are also significantly downregulated in models of progranulin deficiency and are key lysosomal lipids that act as mediators of catabolic processes (12, 14). These data suggest loss of BMPs in progranulin deficient brains may be in part secondary to progranulin-dependent Pld4 deficiency.

We were also interested to test whether the changes we saw in the microglial lysosomal proteome were reflected in microglial RNA-Seq data. To accomplish this, we compared the log2FC’s of *Cx3cr1-Cre+*;*Grn* KO LysoIP samples to *Grn* KO mRNA samples from a previously published microglial RNA-Seq datasets (Figure 4g) (63). 80 of the 111 LysoIP proteins were detected in the RNA-Seq dataset. This included 18 proteins that are significantly altered in the *Cx3cr1-Cre* LysoIP dataset. However, the log2FC values of these proteins did not correlate to the log2FC values of their accompanying transcripts (r^2^ = 0.029). This suggests the majority of the effects of progranulin deficiency on the microglial lysosomal proteome are not secondary to alterations at the mRNA level.

One of the proteins most significantly downregulated in *Grn* KO microglial lysosomes is palmitoyl protein thioesterase 1 (Ppt1). Mutation of this protein leads to neuronal ceroid lipofuscinosis 1 (CLN1), an early onset lysosomal storage disorder with severe symptoms such as microcephaly, hypotonia, and epilepsy (64). Western blot analysis of LysoIP samples from brains of *Cx3cr1-Cre*+, *Syn-Cre*+, and *Ubc-Cre*+ mice demonstrates Ppt1 is effectively absent from microglial lysosomes but not significantly altered in whole-brain or neuronal lysosomes (Figure 4h-k). In our TMT-MS datasets, Ppt1 is significantly down-regulated in *Syn-Cre*+;*Grn* KO LysoIP samples compared to *Grn* WT controls, and while Western blot analysis of these samples does not recapitulate these findings, there is certainly a trend for decreased Ppt1 in *Grn* KO neuronal lysosomes. Regardless, the effect of progranulin deficiency on microglial Ppt1 is much larger than the effect on neuronal Ppt1 in both TMT-MS and western blot analysis.

While Ppt1 is absent in *Grn* KO microglial lysosomes, it is plausible progranulin deficiency leads to accumulation of Ppt1 in non-lysosomal compartments. To test this possibility, we isolated microglia and microglial lysosomes from the left and right hemibrains of 9 to 12-month-old *Cx3cr1-Cre*+ mice, respectively (see Methods). Western blot analysis of both whole microglial and microglial lysosome samples demonstrated significantly decreased levels of Ppt1 in *Grn* KO mice compared to *Grn* WT (Figure 5l-n). This decrease was not significantly different between whole microglia and microglial lysosomes, suggesting progranulin deficiency does not lead to accumulation of missorted lysosomal proteins but instead causes overall decreases of microglial lysosomal proteins.

**Figure 5.**
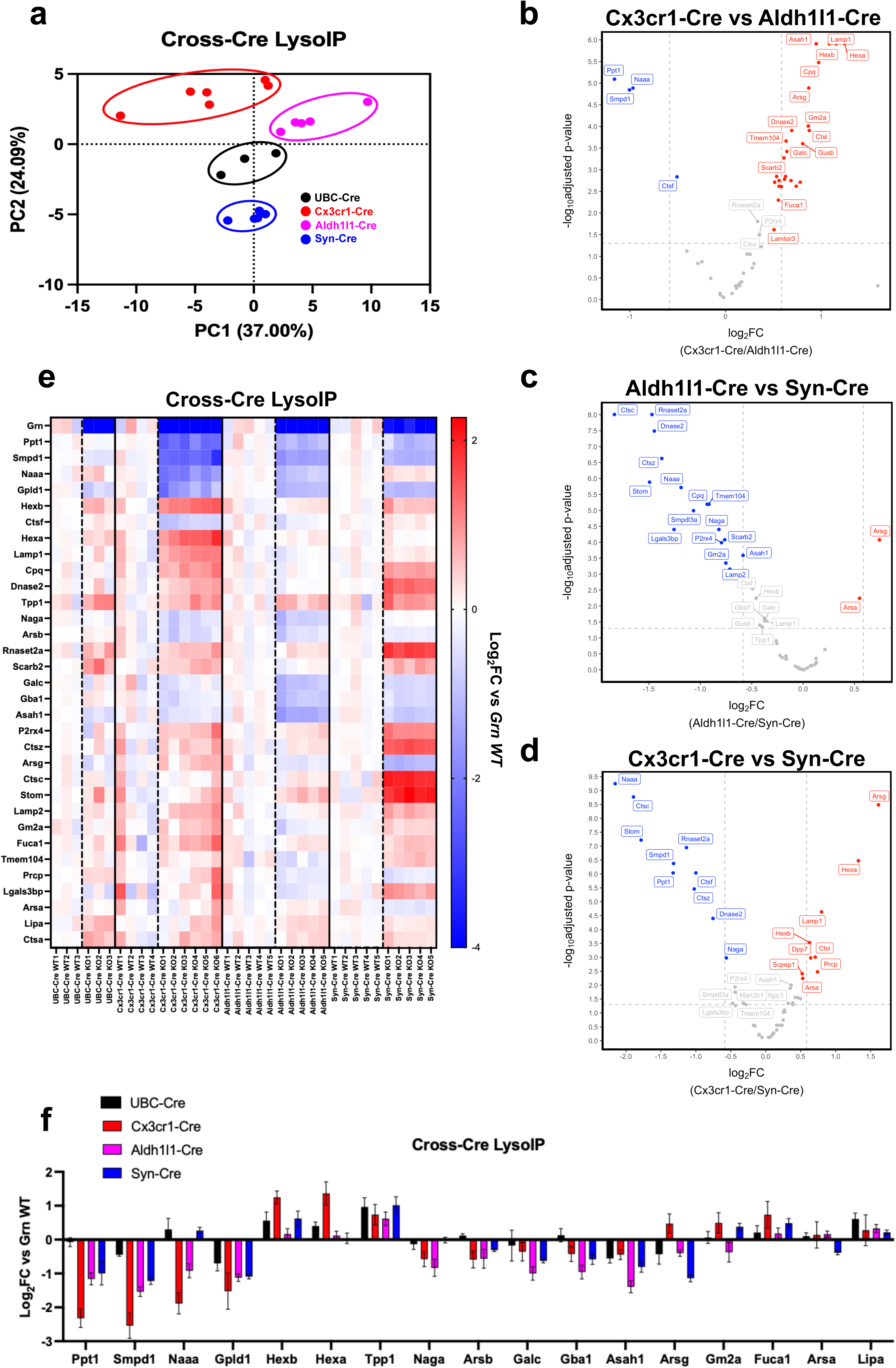
*In vivo* LysoIP datasets demonstrate progranulin-dependent aberrations to the lysosomal proteome are unique across brain cell types. **a** Principal component analysis of *Grn* KO LysoIP log_2_FC values (*Grn* WT vs *Grn* KO) from four separate *Cre* lines. **b** Heatmap depicting log_2_FC values (*Grn* WT vs *Grn* KO) of all proteins detected as significantly altered in *Grn* KO LysoIP samples (*Grn* WT vs *Grn* KO adjusted p-value < 0.05) in at least one *Cre* line. **c-e** Volcano plots comparing log_2_FC values of significantly altered proteins (*Grn* WT vs *Grn* KO adjusted p-value < 0.05) between *Cx3cr1-Cre* and *Aldh1l1-Cre* (**c**), *Aldh1l1-Cre* and *Syn-Cre* (**d**), and *Cx3cr1-Cre* vs *Syn-Cre* (**e**) *Grn* KO LysoIP samples. **f** Bar plot comparing log_2_FC values (*Grn* WT vs *Grn* KO) of key lysosomal proteins detected as significantly altered in *Grn* KO LysoIP samples (*Grn* WT vs *Grn* KO adjusted p-value < 0.05) in at least one *Cre* line.

Progranulin deficiency has also been associated with hyperinflammatory states (24, 65, 66). Further, mutations in several other CLN-related proteins – including Ppt1 – have been shown to increase the activation of the cGAS-STING pathway (67, 68). Similarly, loss of another FTD-related protein C9orf72 also leads to increased cGAS-STING signaling (69). Ppt1 has also been shown to remove the palmitoyl moiety from phosphorylated STING in order to abrogate STING activation (70). To determine if progranulin deficiency led to increased STING signaling similar to other CLN genes, we produced *GRN* KO HeLa cells and measured phosphorylated STING as a function of time after activation with the STING agonist diABZI (71). In the presence of prolonged STING activation, p-STING is known to peak and decline as it becomes digested by the lysosome(72). However, in *GRN* KO cells, STING activation (measured by p-STING levels) was significantly prolonged (Supplemental Figure 4a-d). This suggests progranulin deficiency leads to hyperinflammation in part through prolonged activation of the STING pathway potentially caused by decreased Ppt1-mediated pSTING depalmitoylation. Overall, our data indicate progranulin deficiency impairs microglial lysosomal homeostasis leading to a secondary hyperinflammatory state.

## Discussion

Progranulin deficiency has long been known to affect lysosomal function with loss of progranulin leading to impaired lysosomal size and shape, impaired lysosomal lipid composition, impaired lysosomal pH regulation, and impaired lysosomal protein activity (10, 12-15). Previous work by our lab and others has further shown that progranulin deficiency leads to aberrations of the lysosomal proteome (16, 17, 73). However, the effect of progranulin deficiency on lysosomal composition in the mammalian brain has not been studied in a cell type-dependent manner. To this end, we utilized an immunoprecipitation-based system to isolate lysosomes from specific cell types in *Grn* WT, heterozygous, and KO mouse brains and spleens. TMT-MS analysis of these samples demonstrated that neuronal and microglial lysosomes were most affected by progranulin deficiency; however, progranulin deficiency led to diverging effects between these cell types. When comparing *Grn* KO LysoIP samples, PCA demonstrates clear separation between lysosomes of different cell types (Figure 5a, Supplemental Table 9). Further analysis demonstrates several proteins whose log_2_FC’s between *Grn* WT and *Grn* KO are significantly different between *Cx3cr1-Cre+* and *Aldh1l1-Cre+*, *Aldh1l1-Cre+* and *Syn-Cre+*, and *Cx3cr1-Cre+* and *Syn-Cre+* (Figure 5b-d. Supplemental Table 10) LysoIP samples.

Progranulin deficiency leads to some proteins such as Naaa and Arsg being significantly downregulated in one cell type and unchanged or even upregulated in another; however, comparisons of proteins significantly altered between *Grn* WT and *Grn* KO LysoIP samples in at least one Cre line demonstrate overall trends of most proteins being consistently up- or downregulated between cell types (Figure 5e-f). Despite this, most proteins affected by progranulin deficiency exhibit cell type-specific sensitivities to this deficiency. For instance, Ppt1, while downregulated between *Grn* WT and *Grn* KO LysoIP samples in all cell types, is more downregulated in microglia compared to astrocytes or neurons. In line with these findings, we propose a hypothetical model where certain proteins are either entirely dependent, somewhat dependent, or independent on progranulin for proper localization and/or stability within the lysosome in a cell type-dependent manner (Figure 6). This model is supported by progranulin being found to physically interact with secretory proteins such as the inflammatory mediator sPLA2-IIa as well as the lysosomal proteins Ctsd, Smpd1, and Gba1, the latter two of which are down-regulated in progranulin deficient lysosomes within the mouse brain (11, 74-76). Of note, these protein-protein interaction studies were performed at neutral pH, suggesting they could occur in the less acidic ER or Golgi apparatus.

**Figure 6.**
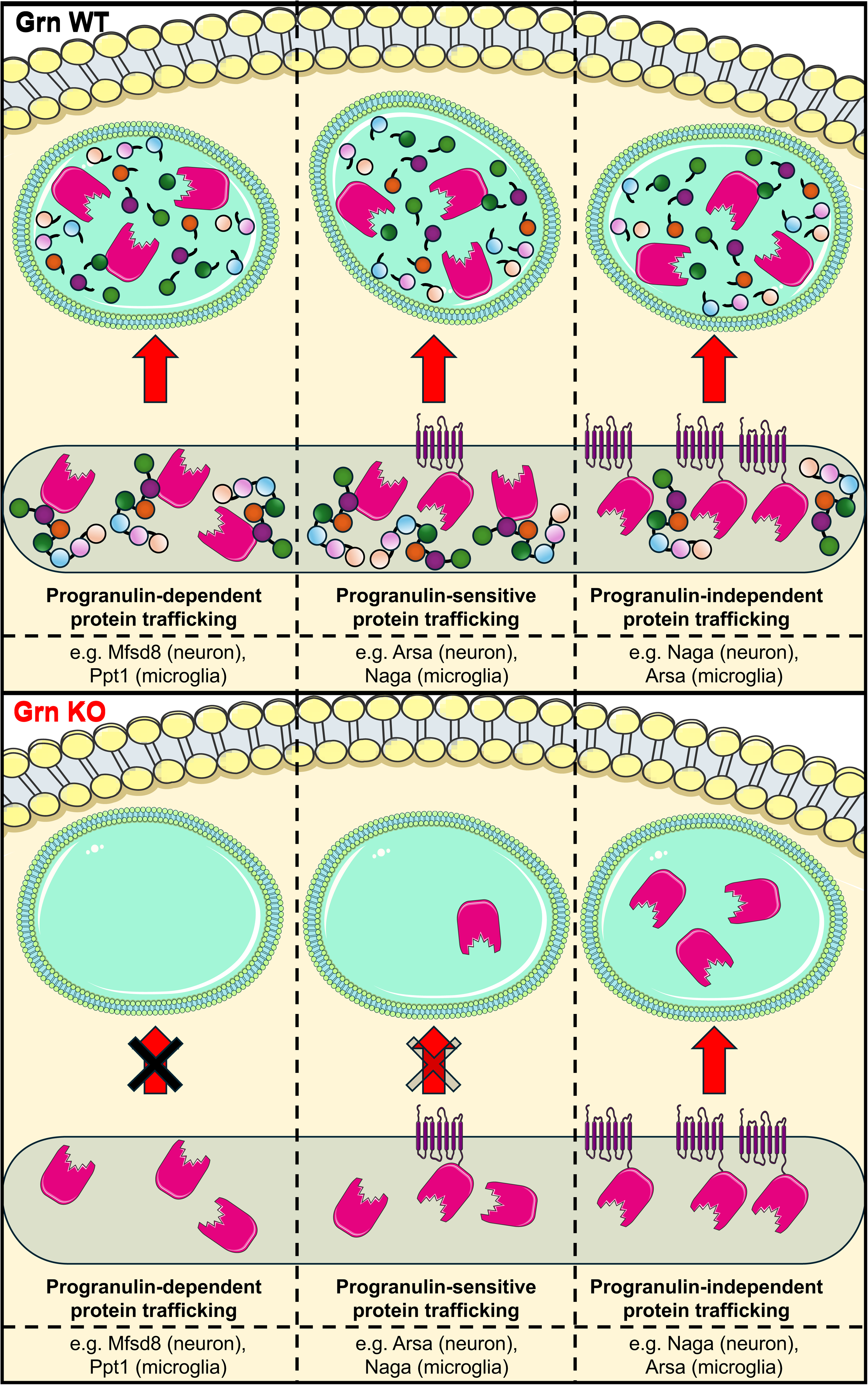
Hypothetical model of how progranulin deficiency potentially alters the lysosomal proteome in a cell-type specific way. Hypothetical model depicting how in certain cell-types, lysosomal proteins may be fully dependent, partially dependent, or independent of progranulin for sorting to the lysosome. The variation between cell types may depend on yet unknown cell-type specific redundant proteins that, like progranulin, also sort lysosomal proteins to the lysosome. Some proteins, such as Mfsd8 in neurons or Ppt1 in microglia, are entirely or nearly entirely dependent upon progranulin for their lysosomal delivery (left panels). Other lysosomal proteins require progranulin for their efficient delivery, but can still complete their journey there, albeit at reduced overall efficiency (middle panels). For other proteins, sorting to the lysosome is undiminished even in the complete absence of progranulin (right panels).

How progranulin affects the lysosomal proteome in a cell type-sensitive manner is not yet understood; however, this could be achieved by composition-dependent multiprotein complex formation or by certain cell types possessing unknown proteins whose functions are redundant to progranulin, while other cell types are entirely dependent on progranulin for lysosomal localization and/or stability of certain lysosomal proteins. This suggests progranulin may function similarly to CLN6 and CLN8, which are responsible for promoting egress of lysosomal proteins from the ER-Golgi complex, or to prosaposin, the precursor for the lysosomal enzyme activating saposins, an established interaction partner for progranulin that is required in precise stoichiometric proportions for efficient transport of progranulin to the lysosome through ternary complexes with Lrp10 or Surf4 (6, 77-79).

Endo-lysosomal dysfunction has been implicated not just in GRN-FTD, but in several neurodegenerative disorders such as Alzheimer’s disease and Parkinson’s disease as well (80). As post-mitotic cells, neurons do not have the ability to dilute impaired lysosomes through cell division. Lysosome membrane repair is also impaired in an age-dependent manner (81). This leads to accumulation of faulty lysosomes in older neurons, impairing proper macromolecule degradation and nutrient signaling. Why some endo-lysosomal defects lead to the accumulation of certain aggregate-prone proteins (e.g., alpha-synuclein, tau, TDP-43, etc.) is yet to be understood, but the correlation between impaired lysosome function and neurodegeneration is well established and may provide a therapeutic avenue for not just GRN-FTD, but for multiple neurodegenerative disorders.

Currently, despite progranulin deficiency leading to 20% of familial FTD, there is no approved treatment for GRN-FTD. Considering the vast network of lysosomal proteins affected by progranulin deficiency in a cell-type sensitive manner, progranulin replacement at the protein synthesis level, and not attempts to correct secondary consequences to lysosomal function or composition, may be the best treatment strategy. Several potential treatments including protein replacement, AAV-technologies, biologics, and small molecule inducers of progranulin expression have emerged in both pre-clinical and clinical trials (14, 16, 82-85). Pathological markers of neurodegenerative diseases appear to accumulate well before the onset of symptoms (86-88), and preventive treatment for these disorders, whenever possible, is likely to provide the best protection against neurodegeneration. Because of its familial inheritance pattern, GRN-FTD is a strong candidate for preventive treatment with progranulin-enhancing agents that aim at increasing the expression of the full-length protein form the remaining functional allele to normalize progranulin levels prior to disease onset. This treatment strategy could potentially stave off the lysosomal aberrations seen in progranulin deficiency and even stall disease onset entirely.

## Materials used in the study

### Antibodies

Mouse anti-Golgin 97 Santa Cruz Cat #: sc-73619

Rat anti-Lamp1 DSHB Clone 1D4B

Rabbit anti-Calreticulin Cell Signaling Cat #: 12238

Rabbit anti-Cathepsin B Cell Signaling Cat #: 31718

Rabbit anti-CoxIV Cell Signaling Cat #: 4850

Rabbit anti-Gm130 Invitrogen Cat #: MA5-35107

Rabbit anti-Calnexin Enzo Cat #: ADI-SPA-860

Rabbit anti-Cathepsin D Abcam Cat #: ab75852

Rabbit anti-Mfsd8 (Cln7) Proteintech Cat #: 24298-1-AP

Rabbit anti-Progranulin Sigma Cat #: HPA008763

Rabbit anti-Ppt1 (Cln1) Proteintech Cat #: 29653-1-AP

Rabbit anti-Psd95 Cell Signaling Cat #: 2507

Rabbit anti-Eaat2 Abcam Cat #: ab41621

Rabbit anti-Iba1 Wako FujiFilm Cat #: 019-19741

Rabbit anti-beta-Actin Abcam Cat #: ab8227

Rabbit anti-Sting Cell Signaling Cat #: 13647

Rabbit anti-phospho-STING (S366) Cell Signaling Cat #: 19781

### Bacterial and virus strains

pX330-U6-Chimeric_BB-CBh-hSpCas9 Addgene Plasmid #: 42230

### Chemicals, peptides, and recombinant proteins

Lipofectamine Thermo Scientific Cat #: 11668030

diabzi-Sting-Agonist-3-(Hydrochloride) Cayman Chemical Cat #: 28054-NC2223805

Rotenone Sigma Cat #: 557368

Sodium Succinate Sigma Cat #: S2378

Antimycin A Thermo Scientific Cat #: J63522.MA

Sodium Ascorbate Sigma Cat #: PHR1279 TMPD

Sigma Cat #: T7394

ADP Sigma Cat #: 01905

Oligomycin Sigma Cat #: O4876

FCCP Sigma Cat #: C2920

Trypsin Thermo Scientific Cat #: 90056

Anti-HA Dynabeads Thermo Scientific Cat # 88836

Anti-Cd11b MicroBeads Miltenyi Biotec Cat #: 130-093-636

### Critical commercial assays

MicroBCA Protein Assay Kit Thermo Scientific Cat #: 23235

NAD/NADH Assay Kit Abcam Cat #: ab65348

### Deposited data

Mass spectrometry proteomics data (Grn WT vs Het vs KO LysoIP raw data)

MassIVE: Dataset accession number: MSV000099828

Password: UTSW2025

ftp://MSV000099828@massive-ftp.ucsd.edu

Experimental models: Cell lines HeLa (89) ATCC Cat #: CCL-2

### Experimental models: Organisms/strains

C57BL/6 (Grn WT, Het, KO) Evers et al. 2017

B6.129S4-Gt(ROSA)26Sortm1(CAG-TMEM192)Dmsa/J Jackson Labs Strain #: 035401

B6.Cg-Ndor1Tg(UBC-cre/ERT2)1Ejb/2J Jackson Labs Strain #: 008085

B6.Cg-Tg(Syn1-cre)671Jxm/J Jackson Labs Strain #: 003966

B6.129P2(C)-Cx3cr1tm2.1(cre/ERT2)Jung/J Jackson Labs Strain #: 020940

B6;FVB-Tg(Aldh1l1-cre/ERT2)1Khakh/J Jackson Labs Strain #: 029655

### Oligonucleotides

Grn KO gRNA: 5’-GGATCGAGCCTTCACGTTGCAGG-3’

Primer: Grn WT For 5’-agtggggctggccacttct-3′

Primer: Grn Mut For 5’-aagattcctcgctgggacatg-3′

Primer: Grn WT/Mut Rev 5’-gaatgctggtgtcagagggcc-3′

Primer: TMEM192-3xHA wt/wt For 5’-CTGGCTTCTGAGGACCG-3’ Laqtom et al. 2023

Primer: TMEM192-3xHA wt/wt Rev 5’-AATCTGTGGGAAGTCTTGTCC-3’ Laqtom et al. 2023

Primer: TMEM192-3xHA fl/fl For 5’-TCCTTGGATATCACCCAGAGT-3’ Laqtom et al. 2023

Primer: TMEM192-3xHA fl/fl Rev 5’-CCACAAAATAACTTTCCCAAGG-3’ Laqtom et al. 2023

Primer: UBC-/Syn-/Aldh1l1-Cre For 5’-GCTGCCACGACCAAGTGACAGCAATG-3’

Primer: UBC-/Syn-/Aldh1l1-Cre Rev 5’- GTAGTTATTCGGATCATCAGCTACAC-3’

Primer: Cx3cr1-Cre WT For 5’- GTGGGAAATCTGTTGGTGGTCCTC-3’

Primer: Cx3cr1-Cre Mut For 5’- GTTAATGACCTGCAGCCAAG-3’

Primer: Cx3cr1-Cre WT/Mut Rev 5’- TAGTCACCCAGACACTCGTTGTCC-3’

### Software and algorithms

Prism GraphPad https://www.graphpad.com/features

ImageJSchneider et al. https://imagej.net/ij/

RStudio Posit Team (2024) http://www.posit.co/Limma

Ritchie et al. DOI: 10.18129/B9.bioc.limma

## Methods

### Cell Culture

All cell culture reagents and plates were purchased from Corning (Corning, NY, USA). Cells were maintained at 8.8% CO2 at 37 °C. HeLa cells were cultured in high glucose Dulbecco’s Modified Eagle’s Medium (DMEM) with 10% FCS and Pen/Strep and supplied from ATCC (CCL-2).

### Generation of *GRN* KO HeLa cells

HeLa cells were transfected with a plasmid containing both Cas9 and sgRNA targeting the R493 region of progranulin as this is one of the most commonly mutated residues(89). Transfection was performed with Lipofectamine 2000 in a ratio of 6:1 (6 μL Lipofectamine: 1 μg DNA). 48-72 hours after transfection, positive cells were selected using puromycin, and individual clones were isolated, grown, and tested for progranulin via western blot.

### STING activation assay

HeLa cells were grown in 6-well plates to 80-90% confluence and were treated with 1 μM diABZI added to 1 mL fresh media for 0, 1, 2, 4, 8, or 12 hours at 37°C. At designated timepoints, cells were rinsed three times with cold PBS on ice, and protein was harvested by scraping cells in 100μL lysis buffer. Cells were incubated with lysis buffer for 15 minutes at 4°C, insoluble material was pelleted by centrifugation at 21,130g for 10 minutes at 4°C, and supernatant was moved to fresh tube. STING activation was measured by western blot analysis using antibodies to human phospho-STING (Ser 366) and total STING.

### Generation of Grn;TMEM192-3xHA fl/fl mice and animal husbandry

The *Grn* mutant mice were originally obtained from the Farese lab and were propagated for at least 50 generations in the Herz lab. Their generation is previously described(13). Animals were maintained on a C57BL/6 J background by heterozygous intercrossing. *Grn* KO mice were bred with TMEM192-3xHA fl/fl mice obtained from Jackson Labs (Strain #035401)(31). *Grn* KO;TMEM192-HA fl/fl mice were then crossbred with either *Ubc-Cre^ERT2^* (Jackson Labs Strain #008085), *Syn-Cre* (Jackson Lab Strain #003966), *Cx3cr1-Cre^ERT2^*(Jackson Labs Strain #020940), or *Aldh1l1-Cre^ERT2^* (Jackson Labs Strain #029655) mice to establish *Grn* WT, heterozygous, or KO; TMEM192-3xHA fl/fl; Cre+ mice. To induce expression of UBC-, Cx3cr1-, and Aldh1l1-Cre, animals were injected at 8 weeks with tamoxifen (120 mg/kg) for up to 5 days. Animals were maintained on a 12 h light/12-h dark cycle and fed a standard rodent chow diet (Diet 7001; Harlan Teklad, Madison, WI) and water ad libitum. No sexual dimorphism of phenotype was observed, and both male and female mice were used for all experiments. All procedures were performed in accordance with the protocols approved by the Institutional Animal Care and Use Committee of the University of Texas Southwestern Medical Center (IACUC APN 2015-101088-G).

### In vivo LysoIP

For isolation of lysosomes from mouse brain, a similar protocol was used as previously described(31, 34). Each LysoIP was individually completed before the next LysoIP was performed. 2-3-month-old (for *Ubc-Cre*, *Cx3cr1-Cre*, and *Syn-Cre*) or 6-month-old (for *Aldh1l1-Cre*) mice were anesthetized with isoflurane and immediately perfused with ice cold PBS (5 mL administered over 4-5 minutes). The brain was removed from the skull, and the olfactory bulbs were removed. Brains were cut into 6 pieces and placed in 950 μL homogenization buffer (PBS + protease inhibitor + phosphatase inhibitor). Brains were homogenized on ice with 25 strokes of plunger, and homogenate was moved to 2 mL microcentrifuge tube. 25 μL of homogenate was moved to separate tube for whole cell lysate. 1 mL of homogenization buffer was added to homogenate to decrease pipetting errors. Insoluble material was spun down at 1000g for 2 min at 4°C. All further steps were performed at 4°C.

1 mL homogenate was added to 100 μL anti-HA Dynabeads which had been pre-washed in 1 mL cold PBS three times before experiment. Homogenate was incubated with Dynabeads with end-over-end rotation for 15 minutes. After incubation, supernatant was removed from Dynabeads, and Dynabeads were washed with 1 mL homogenization buffer three times. After the third wash, Dynabeads were moved to fresh tube. 180 μL lysis buffer was added to Dynabeads and incubated on ice for 15 minutes. Lysis buffer was removed from Dynabeads and moved into fresh tube. For LysoIP of splenic macrophages, the same procedure was used as for the brain, except mice were not perfused with PBS before dissection of the spleen, and 300 μL homogenization buffer was added to homogenate (as opposed to 1 mL with brain LysoIP). All remaining steps were the same as for brain LysoIP.

For whole cell lysate, 200 uL lysis buffer was added to lysate and incubated with end-over-end rotation for 15 minutes. Insoluble material was spun down at 21,130g for 10 min at 4°C. Lysate was moved to fresh tube, and both lysate and LysoIP were stored at -80°C until downstream applications.

### Tandem mass tag mass spectrometry

Sample concentrations were measured by micro-BCA assay and normalized to sample with lowest concentration. Samples were dried for 30 min in a SpeedVac, after which 40 µl of 5% SDS was added to each. Samples were then reduced with TCEP and alkylated with iodoacetamide in the dark. Each sample was loaded onto an S-Trap Micro (Protifi), following which 2 µg of trypsin (Pierce) was added and allowed to digest overnight at 37 °C. Peptides were eluted and dried in a SpeedVac, then were reconstituted in 21 µl of 50 mM TEAB buffer. Samples were then each labeled with 4 µl of TMTpro 16plex reagent (ThermoFisher, Waltham, MA, USA, A44520) and quenched with 2 µl of 5% hydroxylamine and combined in equal peptide amount based on NanoDrop A205 reading. These mixtures were dried in a SpeedVac and reconstituted in 2% acetonitrile, 0.1% TFA buffer.

Peptides were analyzed on a Thermo Orbitrap Eclipse MS system coupled to an Ultimate 3000 RSLC-Nano liquid chromatography system. Samples were injected onto a 75 um i.d., 75-cm long EasySpray column (Thermo) and eluted with a gradient from 0-28% buffer B over 180 min at a flow rate of 250 nL/min. Buffer A contained 2% (v/v) ACN and 0.1% formic acid in water, and buffer B contained 80% (v/v) ACN, 10% (v/v) trifluoroethanol, and 0.1% formic acid in water. at a flow rate of 250 nl/min. Spectra were continuously acquired in a data-dependent manner throughout the gradient, acquiring a full scan in the Orbitrap (at 120,000 resolution with a standard AGC target) followed by MS/MS scans on the most abundant ions in 2.5 s in the ion trap (turbo scan type with an intensity threshold of 5,000, CID collision energy of 35%, standard AGC target, maximum injection time of 35 ms and isolation width of 0.7 m/z). Charge states from 2-6 were included. Dynamic exclusion was enabled with a repeat count of 1, an exclusion duration of 25 s and an exclusion mass width of ± 10 ppm. Real-time search was used for selection of peaks for SPS-MS3 analysis, with searched performed against a list of proteins from the mouse reviewed protein database from UniProt. Up to 2 missed tryptic cleavage was allowed, with carbamidomethylation (+57.0215) of cysteine and TMTpro reagent (+304.2071) of lysine and peptide N-termini used as static modifications and oxidation (+15.9949) of methionine used as a variable modification. MS3 data were collected for up to 10 MS2 peaks which matched to fragments from the real-time peptide search identification, in the Orbitrap at a resolution of 50,000, HCD collision energy of 65% and a scan range of 100–500.

Protein identification and quantification were done using Proteome Discoverer v.3.0 SP1 (Thermo). Raw MS data files were analyzed against the mouse reviewed protein database from UniProt. Both Comet and SequestHT with INFERYS Rescoring were used, with carbamidomethylation (+57.0215) of cysteine and TMTpro reagent (+304.2071) of lysine and peptide N-termini used as static modifications and oxidation (+15.9949) of methionine used as a variable modification. Reporter ion intensities were reported, with further normalization performed by using the total intensity in each channel to correct discrepancies in sample amount in each channel. A co-isolation threshold of 75% and an SPS Mass Matches threshold of 65% was used, and the false-discovery rate (FDR) cutoff was 1% for all peptides.

### Analysis of TMT-MS results

For each TMT-MS run, at least one sample from a Cre- mouse was included as a negative control. All samples were normalized to total protein abundance. LysoIP samples were compared to the Cre- control by one-sample t-test followed by Benjamini-Hochberg procedure with FDR set at 0.05. Proteins were only included in downstream analysis if they were significantly different compared to Cre- control, had an average log_2_FC value > 0.58 compared to Cre- control, and all individual LysoIP sample intensities were greater than no Cre control. Normalized abundances of the Cre- controls were subtracted from the normalized abundances of each sample. Protein quantities were log2 transformed, and the median quantity of protein intensities for each LysoIP sample was subtracted from individual protein intensities to account for differences in cell number and LysoIP efficiency. Differential protein abundance was determined using an ANOVA test applied to the normalized protein quantities using limma(90, 91). Correction for multiple testing was performed using the Benjamini Hochberg method with an FDR set at 0.05. Any protein with adjusted p-value < 0.05 was considered significant. Gene ontology analysis was performed using ShinyGO (92) to compare LysoIP samples and Cre- controls or Metascape to compare *Grn* WT and *Grn* KO LysoIP samples. For ShinyGO, all proteins detected in a single TMT-MS run were used as background proteins to determine enrichment of lysosomal proteins. For Metascape, all 526 proteins detected as LysoIP proteins after filtering were included as background.

### Western blotting

For STING activation assay, cell lysate was first normalized to match lowest concentration as determined by Lowry assay. For all western blotting (except for Mfsd8 immunoblotting), 5x sample buffer was added per sample, and samples were boiled at 95°C for 10 minutes. 10-14 µl per sample was added to pre-cast gel, and electrophoresis was performed at 120V for 70-90 minutes. Samples were transferred to 0.22 µm nitrocellulose membranes using Bio-Rad semi-dry transfer (25V, 1.3-2.5A, 15 minutes). Membranes were blocked with 5% milk in TBS-T and incubated with primary antibody in 3% BSA in TBS-T + 0.02% NaN_3_ either at room temperature for 60 minutes or overnight at 4°C. Membranes were rinsed in TBS-T, and secondary was applied for 60 minutes in 5% milk TBS-T. Membranes were rinsed in TBS-T and either imaged using Licor or ECL.

For Mfsd8 immunoblotting, 5x sample buffer was added to samples and were heated at 37°C for 10 minutes. Samples were run on pre-cast gels as previously described and were transferred to 0.22 µm PVDF membranes using wet transfer apparatus (40V overnight at 4°C). Membranes were blocked with 5% milk in TBS-T and incubated with primary antibody in 3% BSA in TBS-T + 0.02% NaN_3_ overnight at 4°C. Membranes were rinsed in TBS-T, and secondary was applied for 60 minutes in 5% milk TBST. Membranes were rinsed in TBS-T and imaged using ECL.

### Isolation of microglia

Microglia were isolated from *Grn;TMEM192-3xHA fl/fl;Cx3cr1-Cre^ERT2^* was performed on 10-12-month-old mice. Mice were anesthetized with isoflurane and perfused with ice cold PBS (10 mL delivered over 4-5 minutes). Brain was harvested and left and right hemibrains were separated. LysoIP was immediately performed on left hemibrain while right hemibrain was left in dissociation buffer on ice. No more than 4 LysoIPs were performed before microglial isolation was performed. To isolate single cell solution, hemibrains were dissociated by mincing tissue in dissociation buffer on ice until no pieces larger than 1 mm remained. Tissue was moved to 50 mL conical tube and rocked at 37°C for 30 minutes. Dissociated tissue was pressed through pre-wet 70 μm nylon filter using rubber end of a syringe plunger. 2 mL sample preparation media was used to rinse remaining cells through filter. Suspension was centrifuged for at 300g for 10 minutes at 4°C with brake set to low. Supernatant was removed, and cell pellet was resuspended in 4 mL 30% percoll solution. Solution was transferred to 15 mL conical tube and spun at 700g for 10 minutes at room temperature with brake set to zero. Upper myelin layer was removed with 5 mL serologic pipette, and remainder of supernatant was removed. Cells were resuspended in 1 mL sample preparation media, and suspension was moved to fresh tube. Suspension was spun down at 300g for 10 minutes at 4°C with brake set to low. Supernatant was removed.

To isolate microglia, pellet was resuspended in 135 μL MACS buffer and 15 μL CD11b beads and was incubated on ice for 15 minutes. 1 mL of MACS buffer was added, and sample was centrifuged at 300g for 10 minutes at 4°C. 70 μm pre-wet nylon filter was placed on LS columns, and both were attached to MACS magnetic stand. Column was prewet with 3 mL MACS buffer. Supernatant was removed from pellet, and pellet was resuspended in 500 μL MACS buffer. Sample was run over column. Tube was rinsed with 2 mL MACS buffer, and rinse was also run over column. 3 mL MACS buffer was run over column.

Column was removed from magnet, 5 mL MACS buffer was added, and buffer was immediately plunged into fresh tube. This was repeated with another 5 mL MACS buffer. This tube now contained isolated microglia. Cells were spun at 300g for 10 minutes at 4°C with low brake. Supernatant was removed, and pellet was resuspended in 50 μL lysis buffer.

### NAD+/NADH Assay

Procedure is described on Abcam website for kit (Cat# ab65348). In brief, ∼20 mg whole brain from fasted mice was homogenized in 400 μL Extraction Buffer in Dounce homogenizer (40 passes). Samples were centrifuged for 5 minutes 4°C at top speed. Supernatant was passed through a 10 kDa spin column (Millipore Sigma) 30 minutes 4°C at 14,000g to remove NAD-consuming enzymes. Half of sample was reserved as “NAD Total” and half was heated at 60°C for 30 minutes to degrade NAD+ (“NADH”). Added 50 μL sample to 100 μL reaction mix (98 μL NAD cycling buffer + 2 uL NAD cycling enzyme) and incubated at room temp for 5 minutes to convert NAD to NADH. 10 μL of developer solution was added, and colorimetric readout was measured at OD 450 nm at multiple times between 1-4 hours. Samples were compared to 0-100 pmol NAD standards. NAD+ was calculated by subtracting NADH from NAD Total, and NAD+/NADH ratios were calculated for each sample.

### Seahorse Assay

To isolate mitochondria from 12-month-old *Grn* WT and Grn KO mice, mice were anesthetized with isoflurane and perfused with cold PBS (5 mL over 4-5 minutes). Brains were extracted, minced on ice, and placed in ∼10 volumes of isolation buffer. Brains were homogenized by 2-3 slow strokes of a Dounce homogenizer. Sample was moved to fresh tube and spun at 800g for 10 minutes at 4°C. Supernatant was moved to fresh tube and spun at 8000g for 10 minutes at 4°C. Supernatant was removed, pellet was resuspended in 5 mL isolation buffer, and samples were spun at 8000g for 10 minutes for 4°C. Supernatant was removed, and pellet containing mitochondria was resuspended in 100 μL Seahorse buffer. Protein was quantified with BCA, and mitochondria were diluted to a concentration of 100 μg protein/mL. 50 μL (5 μg) mitochondria/sample was transferred to XFe24 Tissue Culture Plate. 4 wells contained 50 μL Seahorse buffer alone as negative controls. Mitochondria were spun down onto plate surface at 3600g for 15 minutes at 4°C. 450 μL of starting buffer was added to each well while Seahorse is calibrated according to manual. Starting buffers and injection port compounds are listed below.

Electron Flow Starting Buffer:

Seahorse Buffer + 10 mM pyruvate + 2 mM malate + 4 µM FCCP

Electron Flow Injection Ports:

Port A: 20 µM (10x) rotenone/injection volume 50 µL

Port B: 100 µM (10x) succinate/injection volume 55 µL

Port C: 40 µM (10x) actimycin A/injection volume 60 µL

Port D: 100mM (10x) ascorbate + 1 mM (10x) TMPD/injection volume 65 µL

Electron Coupling Starting Buffer:

Seahorse Buffer + 10 mM succinate + 2 µM rotenone

Port A: 40 mM (10x) ADP 50 µL

Port B: 20 µM (10x) oligomycin 55 µL

Port C: 40 µM (10x) FCCP/injection volume 60 µL

Port D: 40 µM (10x) antimycin A/injection volume 65 µL

### Quantification and Statistical Analysis

Data analysis was conducted using ImageJ (NIH) and Odyssey Imaging Systems (Li-Cor). GraphPad Prism was used for all statistical analysis except TMT-MS analysis which was performed with the limma package which uses the R programming language. Data are displayed as means ± standard errors (SE). Specific statistical tests including number of samples/condition are specified in individual figure legends.

## Acknowledgments

J. Herz was supported by grants from the NIA (RF1/RF3 AG053391), the NINDS and NIA (R01 NS093382 and R01 NS108115), NIAMS (1R43AR081762) and NINDS (1R43AG084450), BrightFocus (A2013524S & A2016396S), the Bluefield Project to Cure FTD, Harrington Discovery Institute (HDI2019-SI-4479), Alzheimer’s Association (ABA-22-970304), the Presbyterian Village North Foundation, and the Robert J. Kleberg Jr. & Helen C. Kleberg Foundation, Texas Alzheimer’s Research and Care Consortium (TARCC) 1271166 and 1280583. We also thank Tamara Terrones, Brian Emmerson, and Patricia Strackbein for their unrivaled support and brilliant technical assistance.

## Contributions

GCW and JH planned and conceived the project. GCW performed all experiments, analyzed data, and wrote the manuscript. JH edited the manuscript, obtained funding, and oversaw the execution of the project.

## Data Sharing Statement

All raw mass spec data is shared through the MassIVE proteomics database. Data accession number: MSV000099828

All other raw data are shared through the Texas Data Repository.

---

**Supplemental Figure 1.**
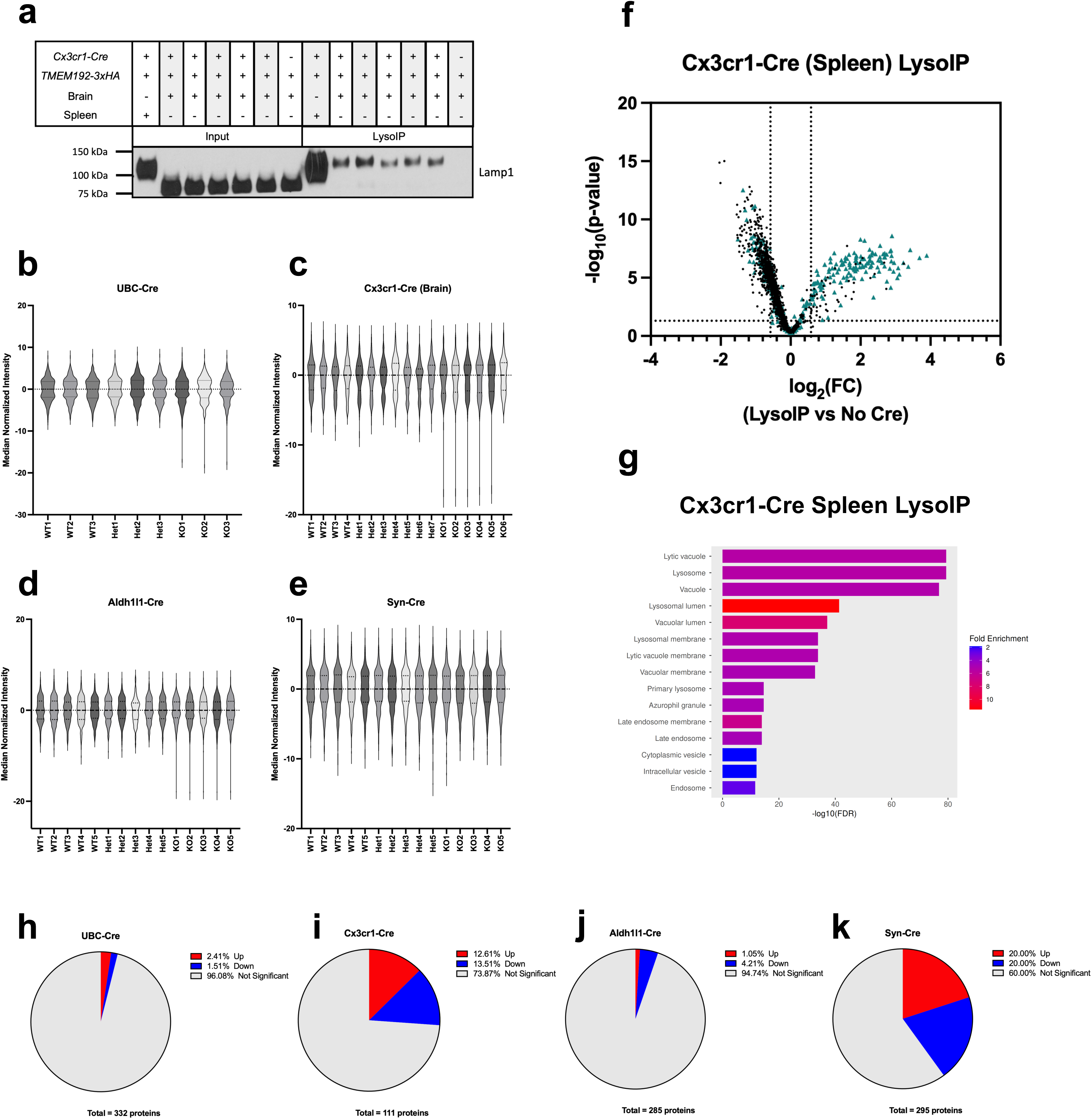
TMT-MS analysis of LysoIP samples demonstrates progranulin-dependent changes primarily in microglia and neurons. **a-d** Violin plots of median-normalized TMT-MS intensity measurements from *Ubc-Cre* (**a**), *Cx3cr1-Cre* (**b**), *Aldh1l1-Cre* (**c**), and *Syn-Cre* (**d**) mice. Gm130 was used as a marker for Golgi apparatus, Lamp1 was used as a marker for lysosomal membrane, calnexin was used as a marker for ER, Cathepsin B was used as a marker for lysosomal lumen, and CoxIV was used as a marker for mitochondria. For *Cx3cr1-Cre*+ samples, cathepsin D was used as a marker for lysosomal lumen due to these samples being imaged on Licor which was not compatible with our cathepsin B antibody. **e** Volcano plot comparing spleen-derived *Cx3cr1-Cre* LysoIP samples to no *Cre* controls (adjusted p-value < 0.05, one-sample t-test followed by Benjamini-Hochberg correction). Known lysosomal proteins are marked as blue triangles. **f** Gene ontology analysis of spleen-derived *Cx3cr1-Cre* LysoIP samples after processing of raw TMT-MS peak intensities. **g** Lamp1 western blot of spleen- and brain-derived *Cx3cr1-Cre* whole lysate, LysoIP, and no *Cre* control samples. **h-k** Pie charts demonstrating percent of proteins significantly up- and down-regulated in whole-brain (**h**), microglial (**i**), astrocyte (**j**), and neuronal (**k**) lysosomes.

**Supplemental Figure 2.**
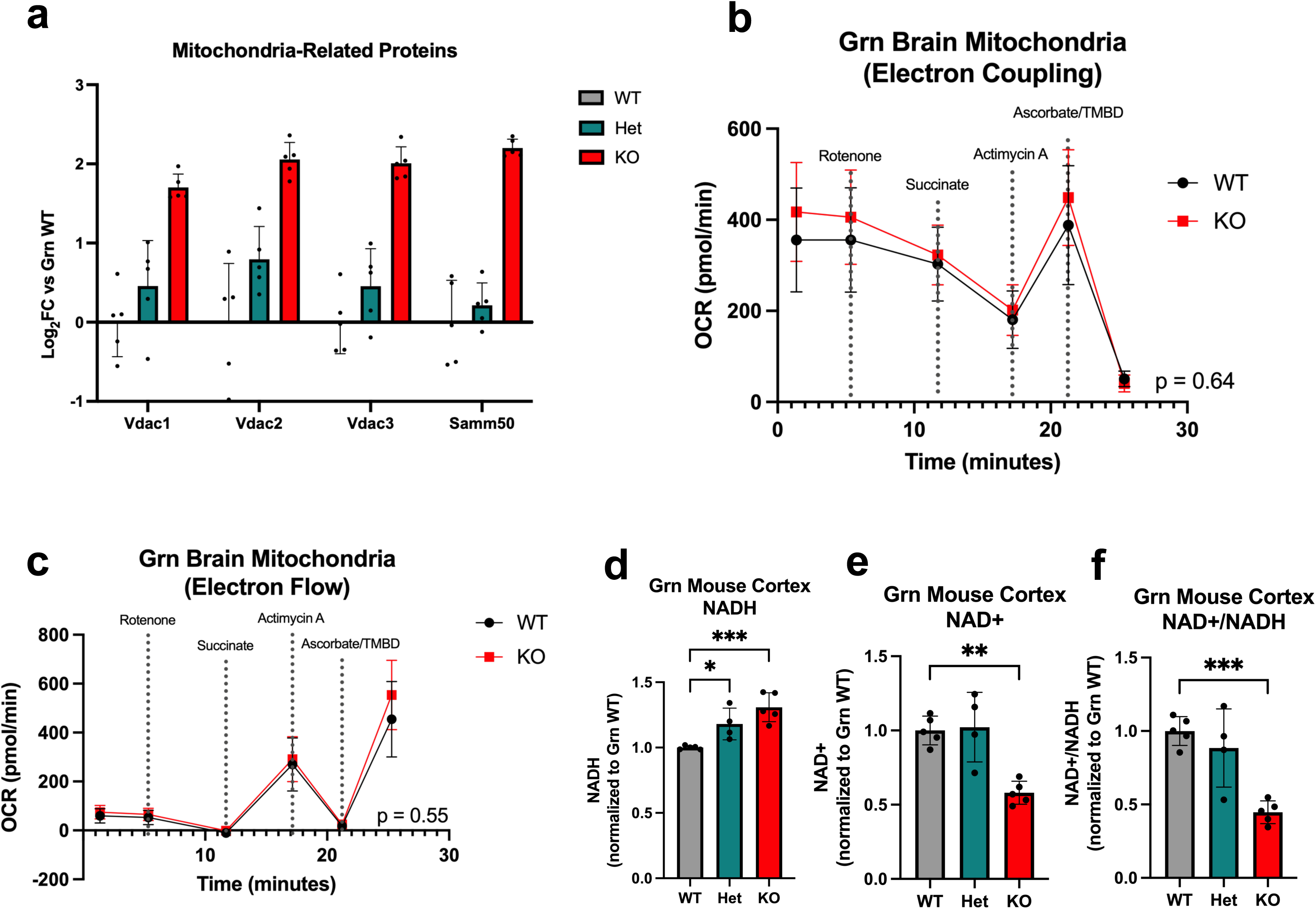
Prgranulin deficiency leads to aberrant mitochondrial protein levels in neuronal lysosomes alongside energetic alterations. **a** Log_2_FC values of mitochondrial proteins significantly increased in *Grn* KO neuronal lysosomes compared to *Grn* WT (adjusted p-value < 0.05). **b-c** Seahorse analysis of electron flow (**b**) and electron coupling (**c**) of isolated mitochondria from 15-month-old *Grn* WT and *Grn* KO mouse brains (n = 3 *Grn* WT and n = 3 *Grn* KO; two-way ANOVA with multiple comparisons and Sidak correction). **d-f** Levels of NADH (**d**), NAD+ (**e**), and NAD+/NADH ratios (**f**) from 15-month-old *Grn* WT, *Grn* Het, and *Grn* KO mouse brains (n = 5 *Grn* WT, n = 4 *Grn* Het, n = 5 *Grn* KO; one-way ANOVA with multiple comparisons and Dunnett correction, *p < 0.05, **p < 0.01, ***p < 0.0001).

**Supplemental Figure 3.**
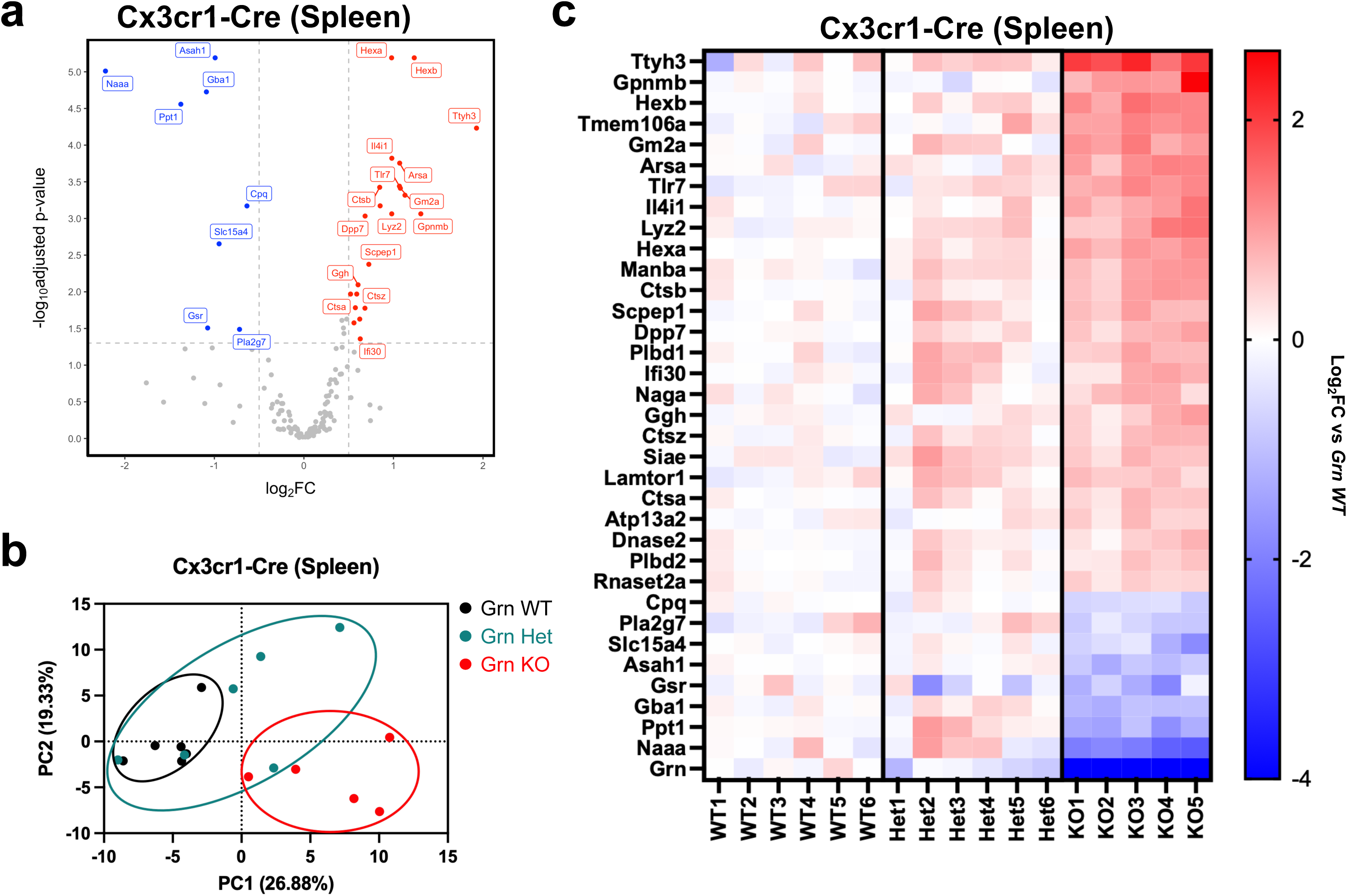
I*n vivo* LysoIP analysis of 3-month-old *Cx3cr1-Cre* mouse spleens demonstrates progranulin-dependent changes in key lysosomal proteins. **a** Volcano plot demonstrating significantly altered proteins (*Grn* WT vs *Grn* KO, adjusted p-value < 0.05) from splenic macrophage LysoIP TMT-MS samples (n = 6 *Grn* WT, n = 5 *Grn* Het, n = 5 *Grn* KO). **b** Principal component analysis comparing log_2_FC values in splenic macrophage lysosomal proteins between *Grn* WT, *Grn* Het, and *Grn* KO mice. **c** Venn diagram comparing significantly altered proteins (*Grn* WT vs *Grn* KO, adjusted p-values < 0.05) in splenic macrophage and microglial lysosomes.

**Supplemental Figure 4.**
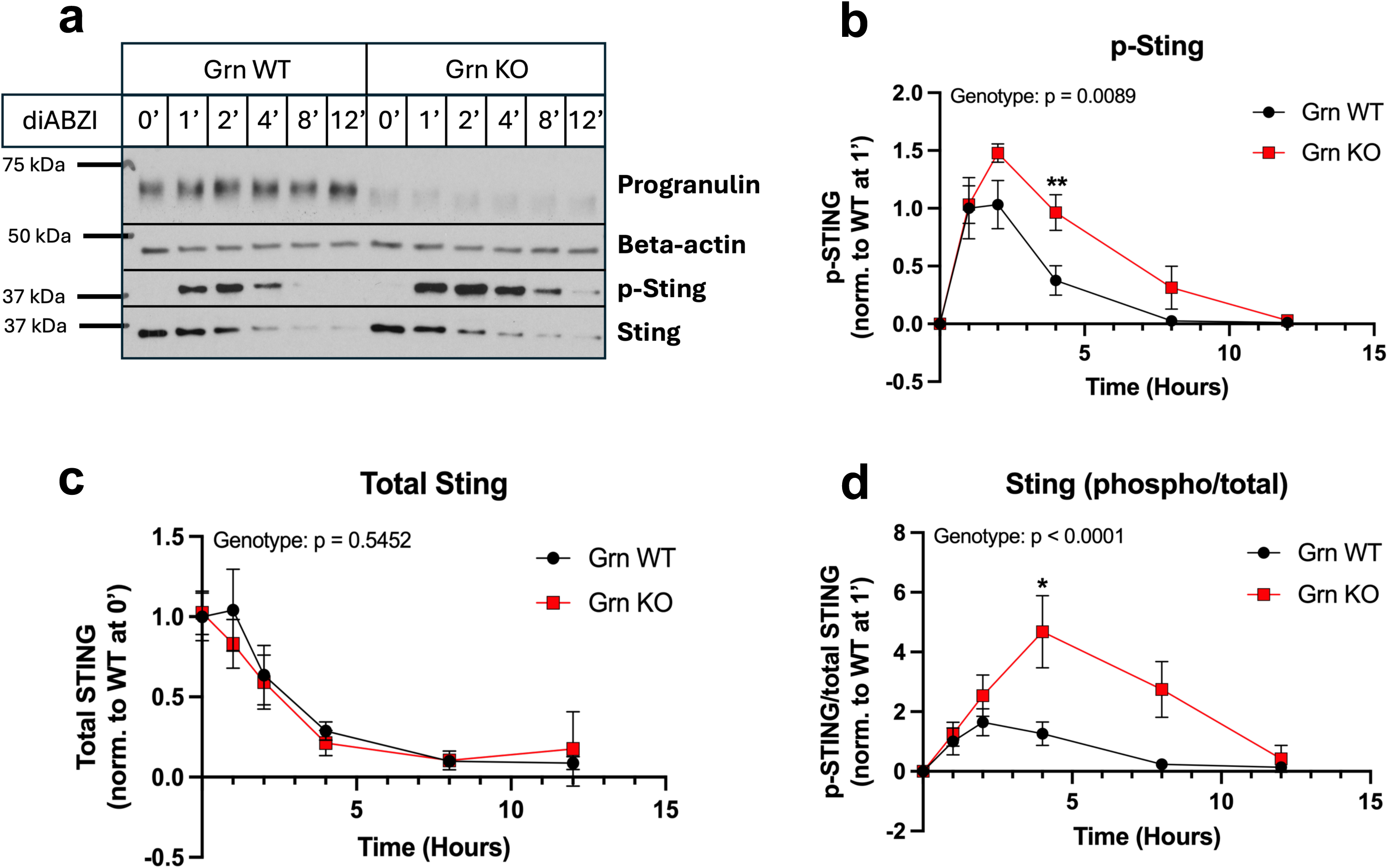
Progranulin deficiency leads to altered STING dynamics *in vitro*. **a** Representative western blot of diABZI-mediated STING activation time course on *GRN* WT and *GRN* KO HeLa cells. Beta-actin is used as a loading control. **b-d** Quantification of p-STING (**b**), total STING (**c**), and phospho/total STING (**d**) of diABZI-mediated STING activation time course (n = 4 *GRN* WT, n = 4 *GRN* KO; two-way ANOVA with multiple comparisons and Sidak correction, *p < 0.05, **p < 0.01).

## Supplemental Tables

**Supplemental Table 1:** Gene ontology analysis of LysoIP proteins significantly altered (*Grn* WT vs *Grn* KO) in at least one Cre line

**Supplemental Table 2:** Log_2_FC and p-values for TMT-MS data for individual mouse LysoIP data from all *Ubc-Cre*+ lines compared to one another

**Supplemental Table 3:** Log_2_FC and p-values for TMT-MS data for individual mouse LysoIP data from all *Aldh1l1-Cre*+ lines compared to one another

**Supplemental Table 4:** Log_2_FC and p-values for TMT-MS data for individual mouse LysoIP data from all *Syn-Cre*+ lines compared to one another

**Supplemental Table 5:** Gene ontology analysis of LysoIP proteins significantly altered (*Grn* WT vs *Grn* KO) in *Syn-Cre*+;*Grn* KO LysoIP samples

**Supplemental Table 6:** Log_2_FC and p-values for TMT-MS data for individual mouse, brain derived LysoIP data from all *Cx3cr1-Cre*+ lines compared to one another

**Supplemental Table 7:** Gene ontology analysis of brain-derived LysoIP proteins significantly altered (*Grn* WT vs *Grn* KO) in *Cx3cr1-Cre*+;*Grn* KO LysoIP samples

**Supplemental Table 8:** Log_2_FC and p-values for TMT-MS data for individual mouse, spleen derived LysoIP data from all *Cx3cr1-Cre*+ lines compared to one another

**Supplemental Table 9:** Log_2_FC and p-values for TMT-MS data for individual mouse, brain-derived LysoIP data from all all Cre lines

**Supplemental Table 10:** Log_2_FC and p-values for TMT-MS data for individual mouse, brain-derived LysoIP data comparing log_2_FC (vs *Grn* WT) values of *Grn* KO samples between Cre lines

